# Application of a Bayesian model based on probability of occurrence to evaluate the effect of an environmental gradient on the turnover of aquatic insect genera in Cerrado streams

**DOI:** 10.1101/2024.10.15.618491

**Authors:** BS Godoy, S Lodi, LG Oliveira

**Author notes:** Corresponding author: Bruno Spacek Godoy, Aquatic Ecology and Fishery program, Amazonian Centre of aquatic ecology and fishery – NEAP, Federal University of Pará - Campus Guamá. Augusto Corrêa street 01, Belém – PA CEP: 66075-110, Brazil.

## Abstract

In ecology, understanding and quantifying the species turnover rates among different locations is essential for a more concise and predictive science. However, little is known about how the turnover occurs among environmentally similar locations and how it may be related to habitat integrity. Using the communities of Ephemoptera, Plecoptera, and Trichoptera sampled from streams in the Brazilian Cerrado, we tested the following hypotheses: a) genus turnover between similar streams will higher in more preserved streams; b) the relationship between proportional genus turnover and the environment is influenced by the taxonomic group. We estimated parameters related to the number of potentials, observed, and alternation of these genera across a habitat integrity gradient in 101 observed streams using a model with Bayesian inference. Genera turnover among environmentally similar sites was directly related to the environmental integrity of the stream. Only the genera Plecoptera showed a pattern similar to the hypothesis we elaborated, showing an increased turnover among well-preserved streams. For the other orders, the turnover presented two peaks in the environmental gradient, one in impacted sites and the other in preserved ones.

## Introduction

A common process in the construction of knowledge in a science is to visit previously debated concepts and adapt recent ideas to the current theory (Vellend, 2010). A current example is rediscovering the importance of the study of species turnover rates and its importance for current theories (Anderson et al., 2011; Veech et al., 2002). Species turnover can be defined as the rate or magnitude of change in species composition along a predefined environmental or spatial gradient (Vellend, 2001). The last decades have been marked by increasing numbers of studies related to differences among communities in different locations (Jurasinski et al., 2009). As a common thread, these studies aimed to understand which factors would structure the communities, such as deterministic (Macarthur and Levins, 1967) or stochastic processes (Jurasinski et al., 2009; Veech et al., 2002; Whittaker, 1957), enabling an adjustment to more current theories (e.g., metacommunities).

The methods used to visualize the species turnover are based on the difference in composition (both qualitative and quantitative) among samples or locations (Anderson et al., 2011; Chao et al., 2005), such as strong and weak points for ecological hypothesis testing. Among the positive points we can highlight the possibility of measuring changes in communities related to environmental gradients. However, these methods do not capture the changes in the communities as a result of the different eco-physiological constraints of the community components in the face of environmental changes (Hutchinson, 1957; Lepš et al., 2006). In addition, the power to explain changes between sites with similar environments is reduced by the limited possibility of separating stochastic effects (such as distinct colonization histories) from environmentally driven effects (Hanski, 1992).

The changes in community components between localities with similar environments have a little explored relevance for ecology, either because the usual methods are poorly documented or because the hypotheses of studies do not aim to distinguish such components (Anderson et al., 2011). Processes such as dispersal, priority effects, or even competitive exclusion can lead to numerical and taxonomic differences at sites with little environmental differences. However, the species turnover rate is limited by the range of organisms that can potentially occur given the environmental conditions shared by the sites (Godoy et al., 2018, 2017; Kraft et al., 2015). When environmental changes are caused by anthropogenic alterations to natural systems, the number of species able to inhabit the site, or local species pool (LSP), is reduced along with the habitat quality (Murphy and Romanuk, 2014). In contrast, local species richness (LSR) across sites tends to be higher among moderately disturbed sites, showing a modal relationship between richness and impact (Huston, 1979; Thorp and Cothran, 1984).

Theoretically, the LSR curve observed at the sites in an environmental gradient is expected to be different from the curve composed of the LSP (Figure 1a). This difference is greater in better preserved sites, indicating a higher species turnover rate among these sites (Figure 1b), which are susceptible to colonization by several species and may already have much of their habitats occupied (Giacomini, 2007). In more preserved sites, the ecophysiological limiting factors are milder, allowing more susceptible organisms to colonize and establish viable populations (Cunha et al., 2020; Godoy et al., 2022).

**Fig. 1.**
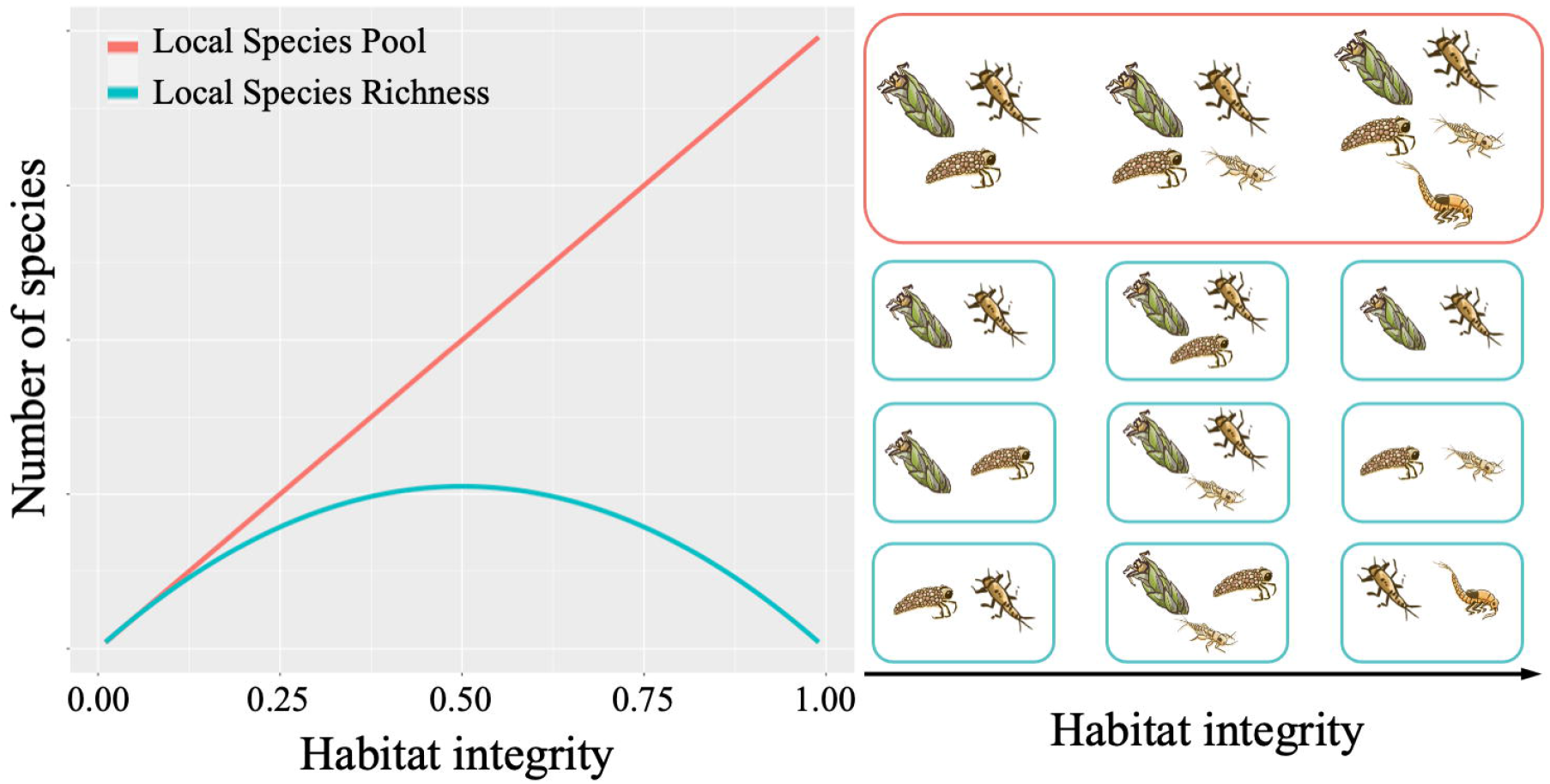
Relationship between species turnover and increased habitat integrity. The larger square represents the local species pool to environment condition, and the smaller ones represent the local species richness.

Although seldom done, observing separately the responses of species to environmental gradients can generate more accurate results and conclusions concerning species behavior in the environment (Godoy et al., 2022; Opedal et al., 2020; Ovaskainen et al., 2017a, 2017b). However, the greatest difficulty for analyses based on operational taxonomic groups is the large number of parameters and relationships to be processed (Giacomini, 2007). In this sense, Bayesian inference applied through hierarchical models becomes an essential tool, making it possible to reduce computational and inferential effort. The Markov Chain Monte Carlo Method (MCMC) has proven to be an efficient methodology for analyses with multiple relations and parameters (Andrade and Kinas, 2008).

The Ephemeroptera, Plecoptera, and Trichoptera (EPT) orders are widely used in studies of stream ecology, especially to study the effects of the environment on communities (Godoy et al., 2022, 2017; Heino et al., 2015b, 2015a, 2013). The main characteristics of these orders are: (1) large number of species; (2) low sampling cost; (3) relationship of the community structure to the environment (Marchant, 2007); and (4) relatively long-life cycle (Céréghino et al., 2003). Using the EPT community, we tested two hypotheses related to the genus turnover among sites with similar environments. The first hypothesis is that genus turnover among environmentally similar streams is directly related to the degree of impact to which they are subjected. We expect more pristine streams to have a higher turnover rate than those that are moderately or severely impacted. As a second hypothesis, we expect that the relationship between turnover and habitat integrity be different depending on the organism group, being higher among Trichoptera due to their increased flight potential compared to Plecoptera and Ephemeroptera.

## Methods

### Study area

The study was conducted in 101 streams of the Rio das Almas River basin, located in the central region of the state of Goiás, Brazil (Figure 2). This river basin presents a gradient from preserved areas to regions with pronounced anthropic impacts (Godoy et al., 2016). Agricultural and cattle raising activities prevail in the impacted areas, with high rates of deforestation and siltation (Strassburg et al., 2017). The study region has a tropical Aw climate and presents a dry period lasting five months (May to September), with average annual temperatures ranging from 24 to 28°C (maximum ranging from 29 to 33°C and minimum from 18 to 22°C). The precipitation in the region varies between 1650 and 1850mm (INMET, 2015).

**Fig. 2.**
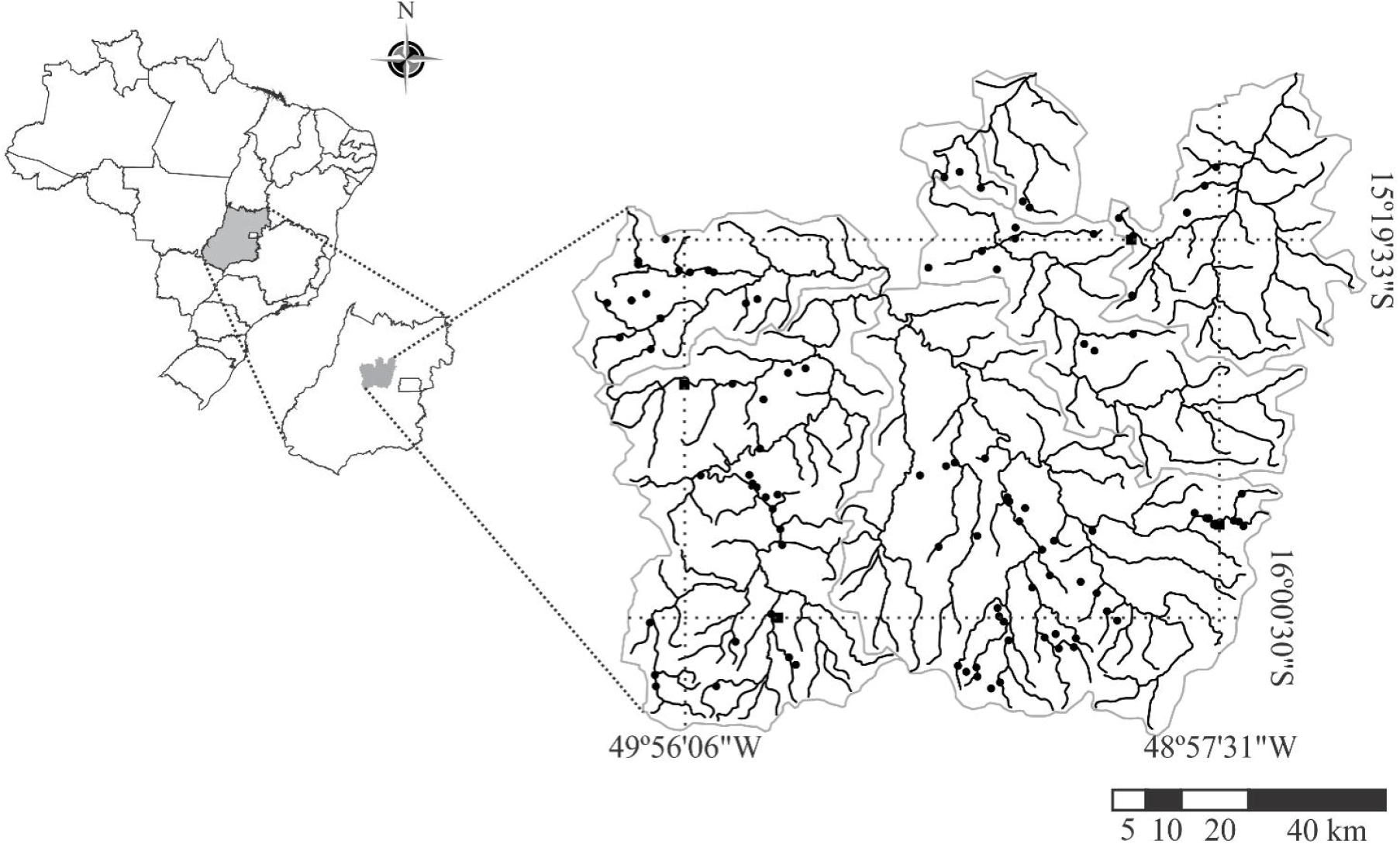
Streams where the community of Ephemeroptera, Plecoptera and Trichoptera were sampled in the Almas River basin from August to October 2008.

### Sampling procedures

We collected insects of the orders Ephemeroptera, Plecoptera and Trichoptera (EPT) from August to October 2008 in four different microhabitats, namely marginal vegetation, stones, bottom litter, and sand. A hand sieve with a 0.025 mm mesh opening was used for 15 minutes for each micro-habitat, amounting to one hour per sampling site. This sampling method was chosen because it yields community composition results similar to those obtained from more exhaustive sampling procedures (Chiasson, 2009).

### Characterization of land use and habitat integrity

We used the Nessimian et al. (2008) protocol to characterize land use and habitat integrity of the areas surrounding and within the streams. This protocol proposes 12 parameters that reflect the environmental conditions of the stream and its surroundings, can be adapted to different biomes, and enables a fast and effective analysis of the environment (Brasil et al., 2020). For the development of this study, we adapted the parameter F1 (see Nessimian et al., 2008), which concerns the width of riparian forest, reducing the classes of vegetation extension. The adaptation was necessary because the Cerrado biome presents naturally less riparian vegetation than the Amazon biome (Ab’Sáber, 2003). According to the proposed protocol, values between 0 and 1 were obtained, where values near 1 indicate reduced habitat changes and near 0 indicate extensive habitat changes. These values constitute the Habitat Integrity Index (HII).

### Data analysis

We used Bayesian inference to test the hypotheses, allowing parameter estimation in a non-linear model with easy fit to the predictions of the study (Andrade and Kinas, 2008; Ellison, 2004; Gelman et al., 2000). We calculated the credibility intervals (CI; 95%) for the parameters of interest using the Markov Chain Monte Carlo (MCMC) methodology, intervals which serve as a direct test of the numerical hypotheses developed (Paulino et al., 2003). The MCMC method is useful to solve complex problems with a large number of parameters, with the advantage that the convergences of its results constitute the *a posteriori* distribution of the hypotheses tested (Ellison, 2004; Gelman et al., 2000). We used 10 independent Markov chains with 10000 iterations sampled at an interval of 50 steps. We used the software R (R Development Core Team, 2020) and the *runjags* package (Denwood, 2016) to perform all analyses, using uninformative prior distributions.

We used a non-linear hierarchical model (see Appendix A for the theoretical development of the relationship) to test the hypothesis that the genus turnover is higher among less impacted streams in the Cerrado biome. The non-linearity of the model is a result of the relationships of the genera occurrence with the environment. The model is based on the difference between the sum of the probabilities of occurrence of all genera given the HII value for the number of genera collected in the streams. The analytical system used to test the hypothesis is:

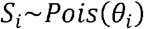

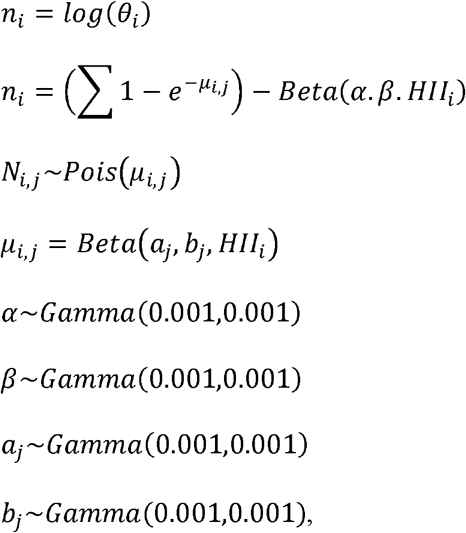

where *S*_*i*_ is the richness observed (LSR) in stream *i, N*_*i,j*_ is the abundance of the genus *j* site *i, Beta*(α, β, *HII*_*i*_) is the beta function of genera turnover for the HII value, and *Beta*(*a*_*j*_, *b*_*j*_, *HII*_*i*_) the beta distribution to the abundance of genus *j* related with habitat integrity. This second function has the parameters *a*_*j*_ and *b*_*j*_ for each genus used in the analysis. The component 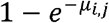 is the probability of occurrence of the genus determined by the HII value in the stream, where the occurrence value is the complement of the probability of no individuals. This probability was estimated using a Poisson distribution with the parameter μ, to modeling the abundance of genus *j* in the site *i*. The sum of 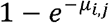 of all genera represent the LSP in site *i*.

We observed the posterior values of the α and β distributions of the beta distribution to test the hypothesis of genera turnover. We observed these parameters in an analysis with all the genera, and in another analysis with the genus grouped by order (EPT). The two parameters are necessary to quantify the turnover relationship and the HII was necessary to display the joint bivariate distribution. Therefore, we estimated the probability of the two parameters being equal to one (when both parameters of a beta distribution are equal to one, the distribution looks uniform). A relationship between alternation and HII that resembles a uniform distribution indicates a lack of effect of habitat integrity on community change.

We also retained the parameter values *a* and *b* of the beta distribution to observe the relationship between the abundance of each genus and the habitat integrity gradient. We categorized the genera into five types of responses to environmental variation, namely Unrelated, Generalist, Sensitive to impacts, Ruderal, and Opportunistic. The categorization was conducted based on the parameters *a* and *b* (Table 1).

**Table 1.**
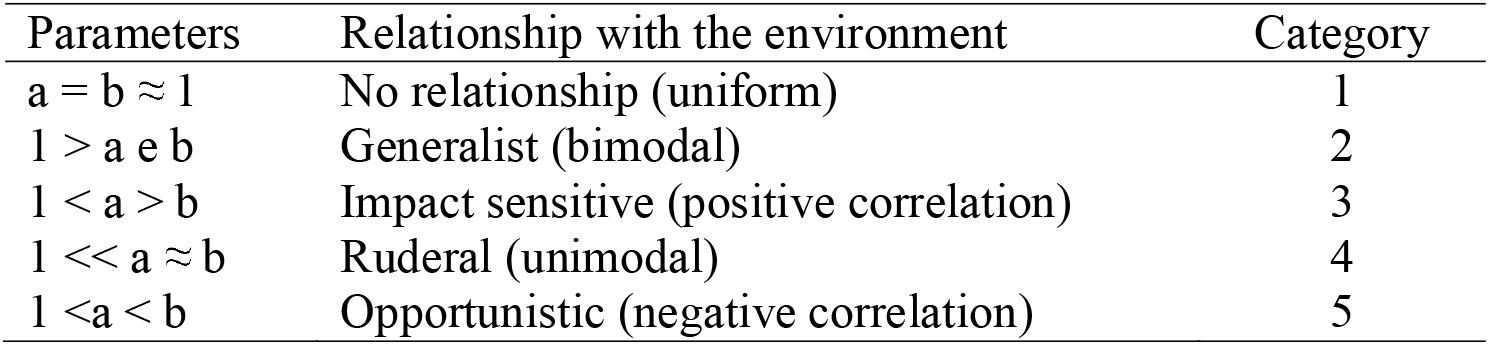
Categories for the distribution of abundances, according to the alpha (a) and beta (b) parameters, for the Ephemeroptera, Plecoptera, and Trichoptera genera.

The grouping provided an estimation of the contribution of each category to the total set of genera within each order. We used a multinomial distribution, since each genus can present only one category, structuring the following model to calculate the *a posteriori* distribution of the ratios:

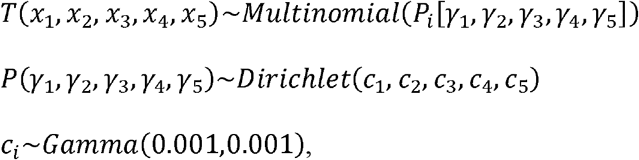

where *T* is the vector with the number of genera in each of the five categories in the order *i*, and *P* the vector of resulting ratios. We repeated this analysis for each order.

We performed the validation of the models in three distinctly moments, the check of model update, the priori sensitive and cross-validation performance (Conn et al., 2018; Kruschke, 2021). The model update was used to identify a possible non-parameter estimation, and it was performed before the burning out of the model using the data. For the priori sensitive we performed the model using uniform priori distribution instead the gamma distribution for the parameter *α, β, a*_*j*_, *b*_*j*_ and compare the results of the models. After the update and sensitive analyses, we used a cross-validation procedure to evaluate the performance of the estimator and the accuracy of the model prediction. In each round of cross-validation, we partitioned the data into two complementary subsets – the training set (with 70% of the observations) and the validation set (with the other 30% of the observations). In each interaction, we removed the validation set and estimated the parameters of the model with the training set. After estimating the parameter, we retained the value of *a*_*j*_ and *b*_*j*_ for each genus and estimated the occurrence of the genus in the validation set, using the value of HII_j_. We used the expected occurrence to calculate the area under curve (AUC) based on the ROC curve of model accuracy and prevalence (Lobo et al., 2008). After completing 1,000 iterations, we used the AUC values calculated in each iteration to estimate the 95% confidence interval, and tested the ability of the model to predict occurrence of genera.

## Results

The aquatic insect abundance ranged from 0 to 279 (93.72 ± 55.48) individuals per stream, and the average richness from 0 to 25 (14.36 ± 5.40) genera per site. We sampled a total of 16,464 individuals which were sorted into 78 genera, being Ephemeroptera represented by 43, Trichoptera by 31, and Plecoptera by 4 genera. The HII showed a large variation, indicating that both preserved and impacted streams were sampled in the region (lowest value = 0.06, highest = 0.95, 1^st^ quartile = 0.38 and 3^rd^ quartile = 0.73).

The turnover of genera was related to the habitat integrity of the streams, either for the whole EPT community or for each order separately. Both the α and β parameters that model the turnover relationship showed a small probability being equal to one in a bivariate distribution (Figure 3). Except for Plecoptera, the *a posteriori* distribution of the CIs of the α and β parameters presented values below one (Table 2). The Plecoptera showed a different parameter estimate (Figure 3c), and thus the relationship between the HII-related genus turnover showed two distinct patterns. The turnover for the components Ephemeroptera, Trichoptera and for the whole community showed two peaks, one at each end of the HII values (Figure 4a, b, d). We should emphasize that the parameters estimated for the entire community and the order Ephemeroptera were similar (Table 2). The number of Ephemeroptera genera is higher in comparison to the other two orders, indicating the importance of this component for the entire community.

**Table 2.** Parameters α and β, and the Credibility Intervals for the turnover of Ephemoroptera, Plecoptera and Trichoptera genera.

| Group | parameter | CI 2.5% | Mean | CI 97.5% |
| --- | --- | --- | --- | --- |
| Entire community | $\alpha$ | 0.13 | 0.14 | 0.16 |
| | $\beta$ | 0.21 | 0.30 | 0.39 |
| Ephemeroptera | $\alpha$ | 0.15 | 0.17 | 0.19 |
| | $\beta$ | 0.34 | 0.78 | 1.90 |
| Plecoptera | $\alpha$ | 1.08 | 2.53 | 4.69 |
| | $\beta$ | 0.35 | 0.78 | 1.90 |
| Trichoptera | $\alpha$ | 0.06 | 0.09 | 0.21 |
| | $\beta$ | 0.07 | 0.22 | 0.42 |

**Fig. 3.**
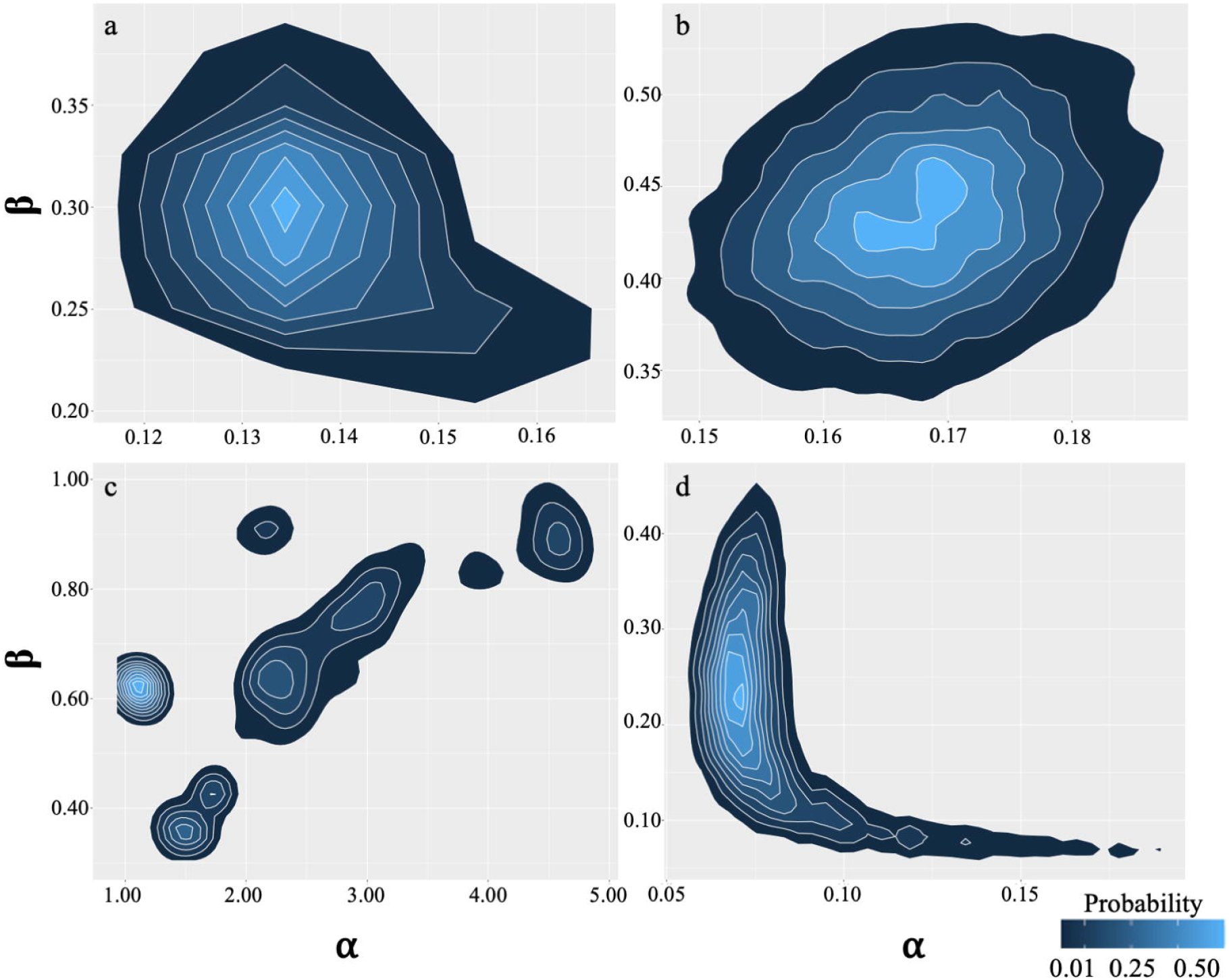
Joint a posteriori distribution for the α and β parameters for genus turnover. a) entire community, b) Ephemeroptera, c) Plecoptera, and d) Trichoptera.

**Fig. 4.**
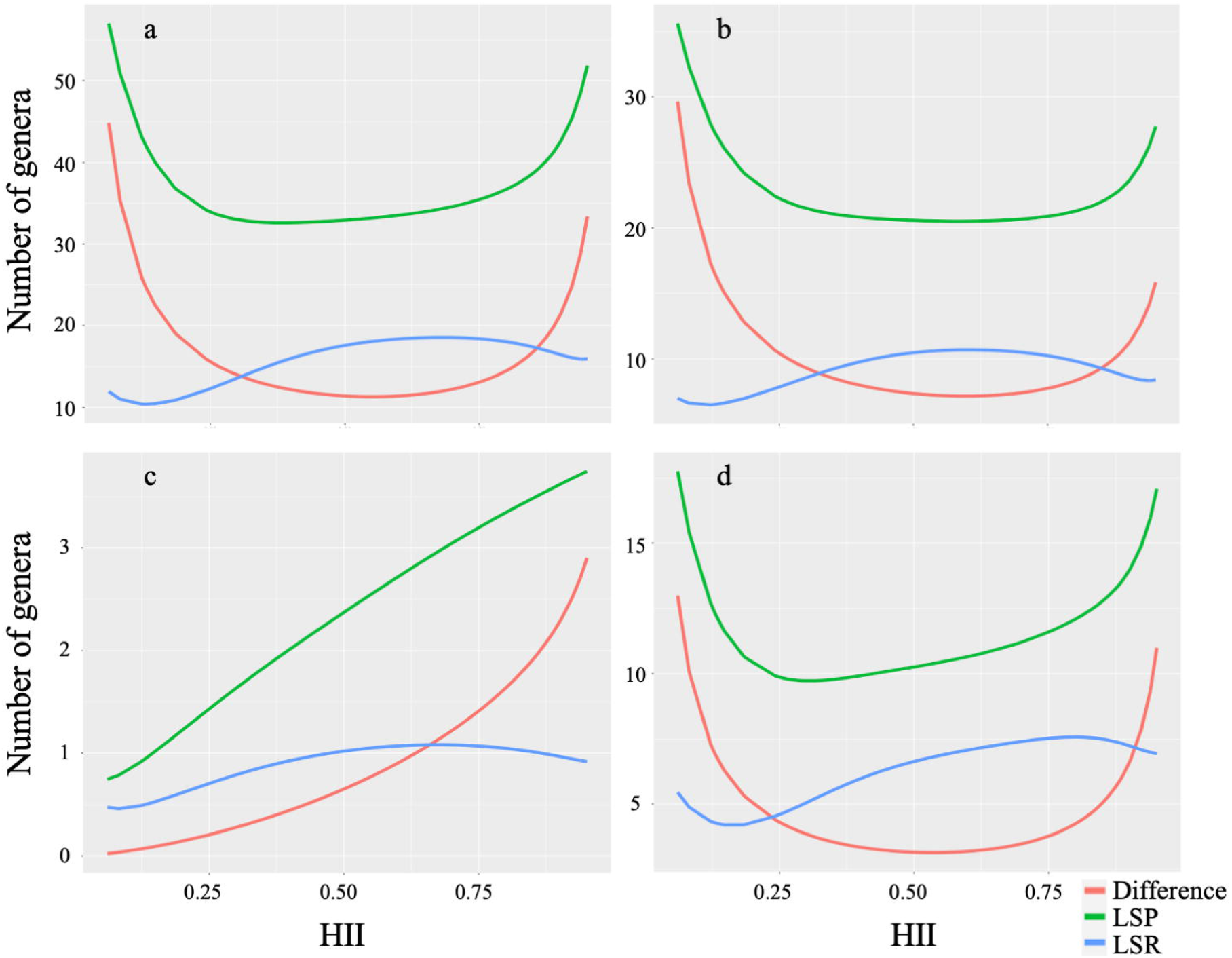
Relationship between the habitat integrity index and the LSR, LSP, and the difference of these two values (diversity turnover among sites). a) entire community, b) Ephemeroptera, c) Plecoptera, d) Trichoptera.

For the order Plecoptera the α and β values showed great dispersion compared to the other community components. However, despite this high dispersion, the probability density is small for values far from the mean (Figure 3c), thus enabling analyses with these values. The result for Plecoptera indicated a higher alternation value for more preserved sites (Figure 4c). The behavior of the order Plecoptera was identical to what we expected according to our hypothesis, which did not occur for the other components of the community tested.

Most genera showed a larger population size in more preserved sites (Table 3). The category of impact-sensitive genera had the highest proportion in all orders (Figure 5). All genera were related with the HII, showing that the index can capture relevant elements of the environment that affect organisms. The categories of genus relationship to the environment within the orders Ephemeroptera and Trichoptera had similar proportions (Table 3 and Figure 5). However, we can point out a subtle difference, whereby the order Trichoptera had a higher proportion of generalist genera, while Ephemeroptera had many opportunistic genera (Figure 5). The high number of genera in groups other than the impact-sensitive category may explain the observed pattern of genus turnover for these two orders (Figure 4b, d). Finally, all genera within the order Plecoptera were positively related to the abundance and HII values.

**Table 3.**
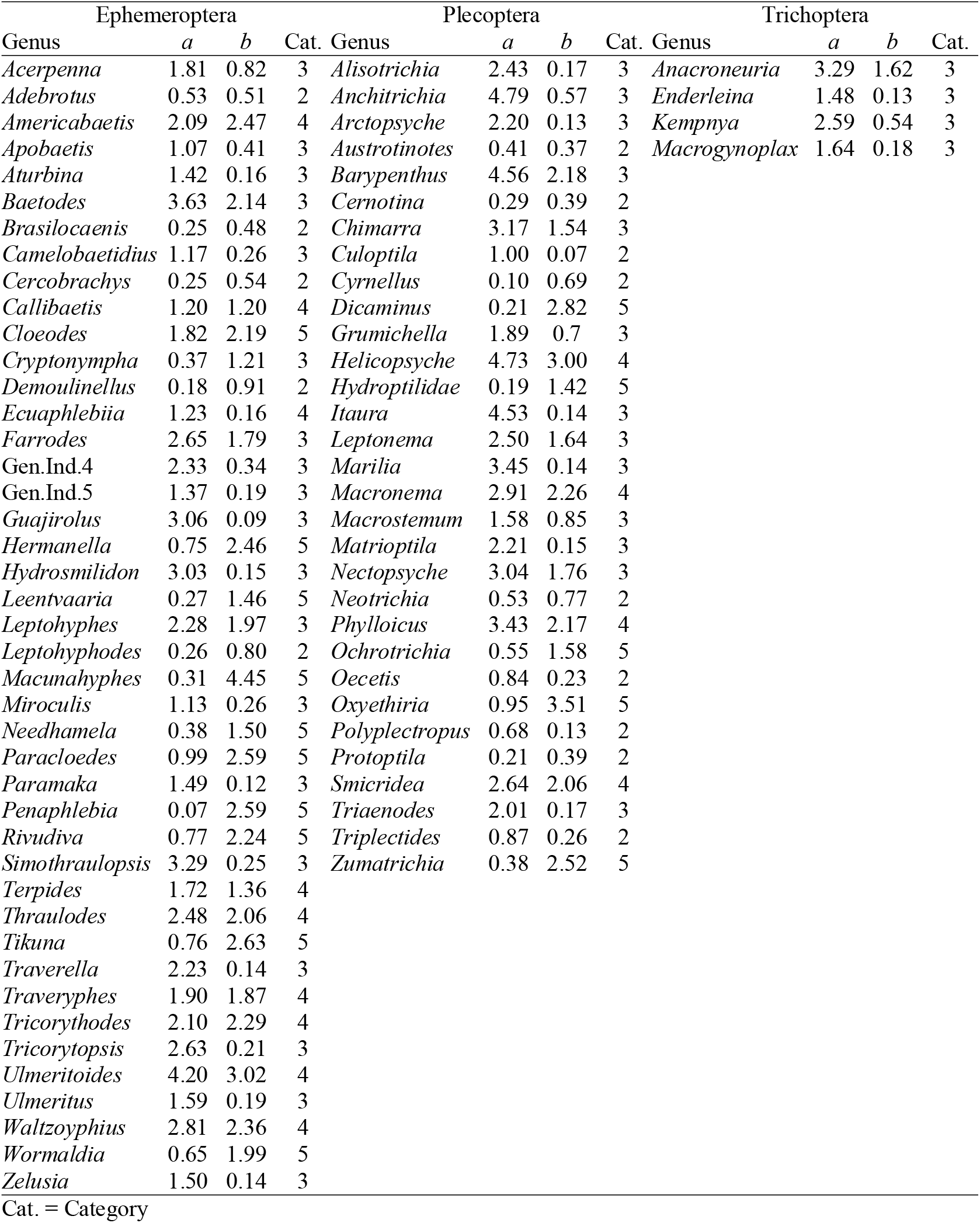
Parameters *a* and *b* for the environment-related distribution of abundances for the genera Ephemeroptera, Plecoptera and Trichoptera. Cat. = category as defined in Table 1.

**Fig. 5.**
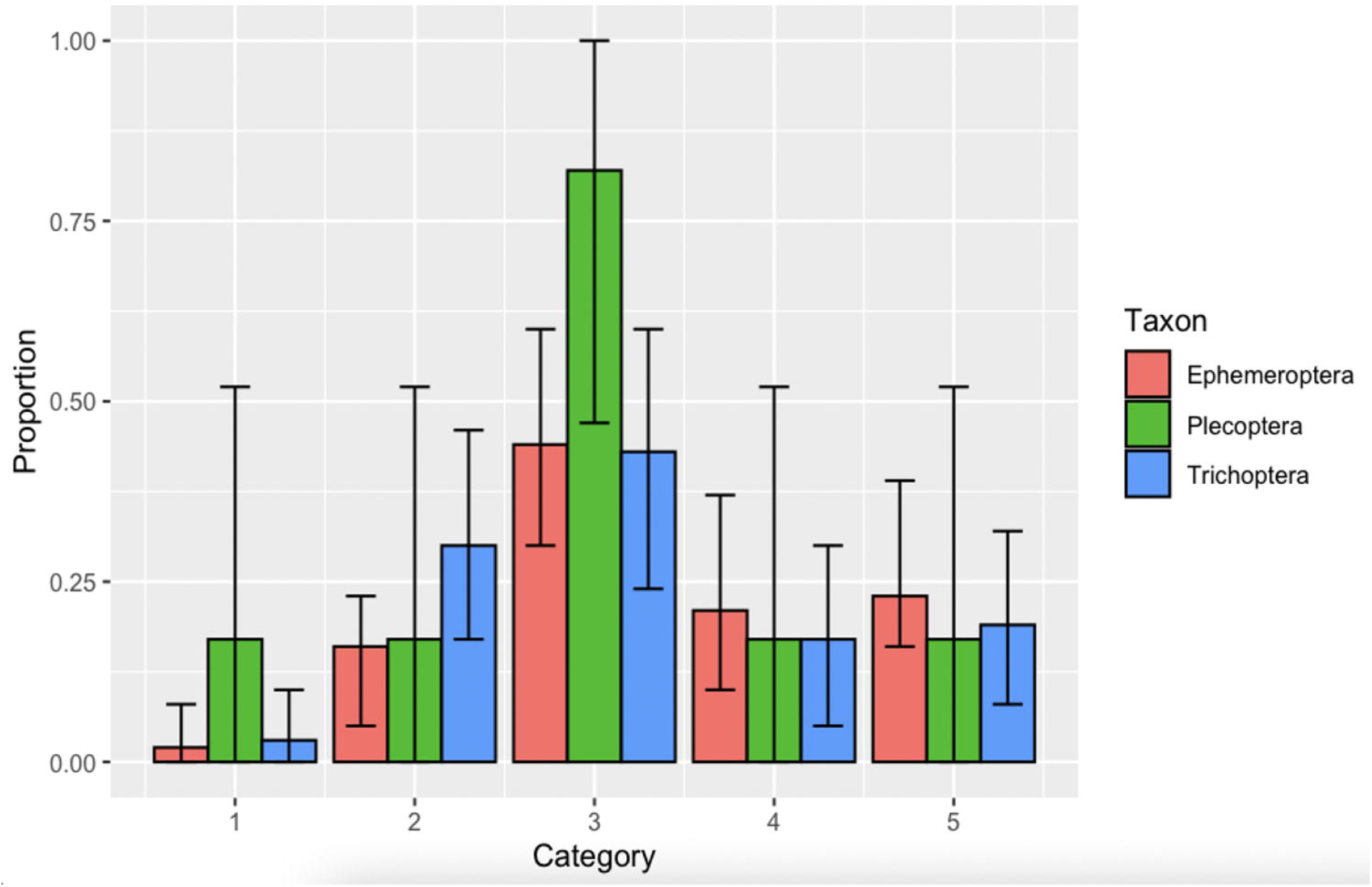
Proportion of genera distributed across the five categories of relationship of abundance to habitat integrity for the orders Ephemeroptera, Plecoptera, and Trichoptera. The error bar represents the 95% credibility interval.

The models were validated for all three tests (model update, priori sensitive and cross-validation). When we changed the priori distribution from a uniform to a gamma, the posteriori distribution did not show a relevant difference (Appendix B). The AUC statistic for the 1,000 iterations was higher than the 0.5 threshold established to validate the model (AUC mean = 0.74, with the 95% CI ranging from 0.62 to 0.81). Lower AUC values were recorded for the order Thricoptera (AUC = 0.61), and the order Plecoptera had the highest AUC values (0.83).

## Discussion

The genus turnover had a complex behavior concerning IIH, making it necessary to understand the context of the study area. A large part of the study area was modified by agricultural practices, where most of the streams were located in an impacted matrix (Colman et al., 2022; Strassburg et al., 2017). The main impact sources of agricultural activities are the removal of riparian forest and siltation, and not pollutant emissions.

The Arrenius Relationship (Preston, 1960) predicts a reduction in the number of taxonomic types when we reduce the area. Thus, we would expect a greater number of genera to occur in degraded sites than in preserved sites, solely due to the area effect. Given that the proposed model calculates the potential occurrence of the genera, the higher regional richness of impact-tolerant organisms would increase the potential richness for this group. However, this increase would be the indirect result of a larger pool of impacted streams, rather than a larger pool of genera acclimatized to adverse conditions.

The number of potential genera in the study was increased twice – in impacted sites due to the area effect and in well preserved sites where the environmental conditions are less adverse enabling the survival of a higher number of species. However, the observed richness has a bell-shaped relationship with HII, with higher values in moderately impacted sites, according to the index used to measure environmental modification. The result is similar to that found in streams in New Zealand, where impacts are caused by flooding, increased flow (Townsend et al., 1997; Townsend et al., 1997), and habitat instability (Death and Winterbourn, 1995). In the New Zealand studies the disturbances were discrete temporally, unlike in this study, where the adversity faced by the streams is constant.

Despite the results of our study showing a higher richness in sites with intermediate impact levels, we should be careful when comparing them with the predictions of the intermediate disturbance model (Giacomini, 2007; Tokeshi, 2009). The reduction in interactive forces among species due to a disturbance would not necessarily lead to reduced rates of competitive exclusion (Chesson, 2000; Chesson and Huntly, 1997), refuting the main argument of the intermediate disturbance hypothesis. However, the models tested for these conclusions are purely theoretical and spatially implicit, such that the effect of the spatial arrangement of communities and dispersal do not affect community structure. Mass effect and the spatial configuration of communities are common elements in ecological systems (Godoy et al., 2022, 2019, 2017; Robson and Chester, 1999), allowing sites with higher taxonomic richness and/or abundance to supply individuals to sites with lower diversity (Hanski and Ovaskainen, 2000; Kunin, 1998). Thus, sites with intermediate disturbance levels may present a higher richness because they are receiving propagules from both impacted and preserved sites.

The turnover of species among different sites caused by both environmental and historical factors has been discussed extensively in ecology (Anderson et al., 2011; Jurasinski et al., 2009; Lu, 2021; Veech et al., 2002). Although it has become commonplace in ecology, the great challenge is no longer the use and comparison of different methods to examine the observed patterns, but rather the development of robust mechanisms able to identify the key processes (Anderson et al., 2011). The phenomenon may only be fully understood when the mechanisms used to analyze the joint abundances of the taxonomic groups explored can be used as tools to explain the mechanisms that structure the diversity changes (Tuomisto, 2010a, 2010b).

The model presented in our study has the advantage of enabling the direct observation of the relationships between the genera and the environmental gradient, allowing the understanding of how the turnover process occurs in different locations, as well as explaining their complex behavior. The coefficients of the Beta distribution for each genus show a large number of genera sensitive to impacts to the stream habitat.

However, approximately 20% of the genera showed an increased abundance in sites with impacted habitat, and 17% of the genera presented abundance peaks in sites with intermediate habitat integrity. Such a pattern of proportions of response groups affects the turnover of genera among sites with different degrees of impact, indicating the existence of specific fauna for each type of environmental modification.

Genera tolerant to increased environmental adversity were found in the orders Ephemeroptera and Trichoptera, with no representatives in Plecoptera. The order Plecoptera also showed the least amount of morphological and taxonomic types among aquatic insects in the Neotropical region (Tomanova et al., 2006; Vinson and Hawkins, 2003). In addition, its species usually require cold and well-oxygenated running waters, limiting the range of occurrence (Braun et al., 2014; Resh and Rosenberg, 2010; Silva et al., 2018). The restricted degree of tolerance added to a low taxonomic and functional diversity causes the Plecoptera genera to have a strong relationship with habitat integrity.

The model tested in this paper presents interesting and essential characteristics for studies on diversity turnover among sites. The main advantage is the direct relationship between the ecological or physiological constraints of taxonomic groups and the turnover rates across an environmental gradient. The parameters estimated by the model are direct responses to fluctuations in the population sizes of the stream genera, directly reflecting the processes determining the diversity turnover. The curves for the quantity of observed, potential, and differential genera were generated explicitly by ecological processes, directly answering the questions posed in the paper.

The model presents two main drawbacks, one relative to the time required for the MCMC iterations, and the other to the non-inclusion of space as a determining element in the processes studied. The relatively large time required for parameter estimation using this method has already been reported (Sturtz et al., 2005), especially when the number of parameters is high. Despite this limitation, the time expenditure is small compared to the amount of information extracted on the genera turnover patterns and on the natural history of the groups studied. The fact that the model is not spatially explicit makes it difficult for us to derive information on the role that spatial dispersion or even spatial autocorrelation may play in the turnover of genera conditioned to habitat integrity. A step forward to better understand these turnover patterns will be to include space as a covariate affecting the local abundances of the genera studied.

Our results present an intriguing and at the same time worrisome scenario concerning the conservation of aquatic insect diversity in the Cerrado region. The number of well-conserved streams is becoming smaller and smaller on a global scale (He et al., 2019; Reid et al., 2019; Vörösmarty et al., 2010), and the Cerrado is no exception (Colman et al., 2022). However, the number of genera that a well-conserved stream can exhibit both potentially and observed is greater than the number that impacted streams may present. Thus, the delimitation of practices that aim to maintain the integrity of habitats in the Cerrado streams becomes urgent. This measure, in addition to improving the streams that are already targets of anthropic activities, enables the local and regional diversity of aquatic insects to be maintained.

## Supporting information

Mathematical framework and algorithm for the calculation of the potential and observed richness

## Aknowledgments

We thank J Simião-Ferreira, LL Queiroz, LM Camargos for field assistance. This study was supported by research grants from CNPq (process 303835/2009-5 and 475355/2007-5) and UFPA (process 02/2022, PAPQ/PROPESP).

## Data availability statement

Data available at https://doi.org/10.5281/zenodo.8329240.

