## Supplementary material for "Application of a Bayesian model based on probability of occurrence to evaluate the effect of an environmental gradient on the turnover of aquatic insect genera in Cerrado streams": Mathematical framework and algorithm for the calculation of the potential and observed richness

Ecology and Evolution

Godoy BS, Lodi S, Oliveira LG

**Appendix A**

Mathematical framework and algorithm for the calculation of the potential and observed richness

According to the multidimensional niche concept the distribution of species is a response to eco-physiological conditions in an n-dimensional space, where each axis represents a resource or condition. The species present in a community would be related to deterministic processes (May and Arthur, 1972; Tuomisto and Ruokolaine, 2006), being a general law and an axiom for deductive theories. Each species has a unique environmental requirement, its occurrence can be described using the function:

(1) $o\left( s_{i} \right)=\frac{e^{\left( x_{1}\beta_{1}+x_{2}\beta_{2}+\cdots+x_{n}\beta_{n} \right)}}{e^{\left( x_{1}\beta_{1}+x_{2}\beta_{2}+\cdots+x_{n}\beta_{n} \right)}+1}$,

where o(si) is the occurrence of species *i*, (*x*_1_,*x*_2_,...,*x*_n_) represents the *n* environmental variables of the site and (*β*_1_, *β*_2_,...,*β*_n_) the eco-physiological limitations that species *i* can withstand.

We assume that *O* = (*o*_1_,*o*_1_,...,*o*_m_ ) is the family that comprises the functions of occurrence of *m* species in a given type of environment, and that the species have different environmental requirements in at least one dimension (*o*_i_ ≠ *o*_j_, where *i* > *j* and *i*, *j* = 1,2,...,*m*). Therefore, when the environmental variables change, the responses will differ among the species, which leads to a species turnover through environmental change. Thus, we can observe a *continuum* of associated fauna that alternate following an integrity gradient. The environmental integrity of the aquatic system is defined here as the physical conditions of the water bodies, such as the riparian vegetation, sediment type, and other hydromorphological characteristics.

The distribution of species would then follow two hierarchical processes: a) locally, where the determining factor would be the interaction of the organism with the environment, and b) among sites, or regionally, where demographic processes would act decisively. Dispersal processes would enable species to arrive on site, while habitat selection processes or biotic interactions would act as filters for which species would establish themselves on site (Godoy et al., 2022, 2017; Heino and Paavola, 2003). Until now, we don’t show the mechanism to able the persistence of multiple species with same environmental requirements. The most relevant argument to explain the coexistence of species with similar requirements it the *d* function of niche overlap between different species (May and Arthur, 1972).

Environmental adversity is related to the group (*x*_1_,*x*_2_,...,*x*_n_), where its increase, as a result of reduced integrity, would lead to a reduced amount of species supported by the environment, due to the limitations established by (*β*_1_, *β*_2_,...,*β*_n_). The reduction of this adversity, however, leads to an increased number of individuals in each species due to *φ*_i_. In a model where *d* have no influence with the coexistence of species, showing no relationship in species interactions with environmental occupancy, we have a linear increase in species richness with increasing environmental integrity (Figure A.1).

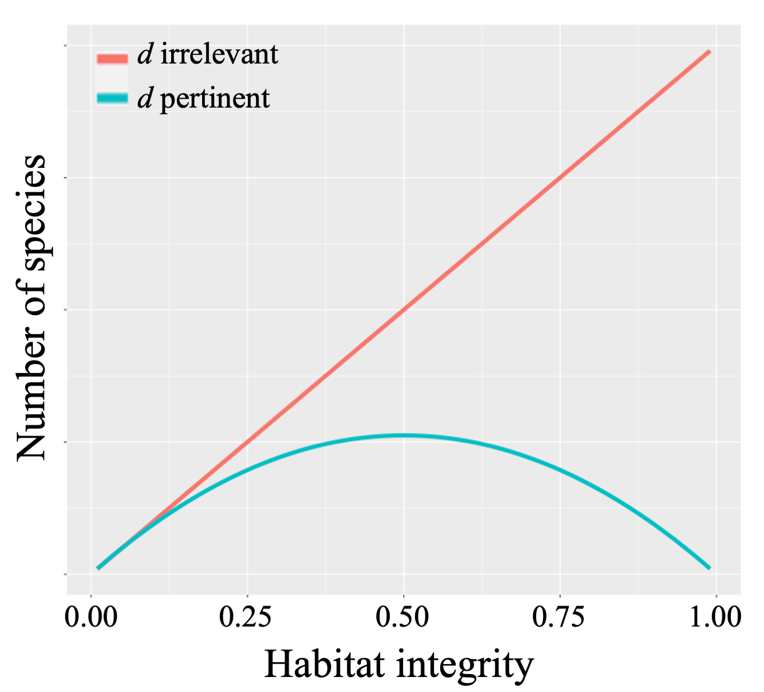

Figure A.1. Relationship between species turnover and increased habitat integrity related with the importance of *d*.

In contrast, when *d* is pertinent to the occupancy of species, in sites with high integrity, the function related to the increased number of individuals (*φ*_i_) reaches its peak value. Therefore, even if there is a large number of species able to colonize the site, the interactions among the organisms becomes a major factor in the number of species. Keeping *total of equivalent individuals in a community* (Hutchinson, 1957) constant for analytical purposes, such increase in individuals per species can only be supported by reducing the number of species per site (Figure SI).

Joining the estimated richness curves related with *d*, we can observe a surplus of species able to inhabit more preserved sites, when compared to the number found on the site. Thus, *the turnover of species in more preserved locations is higher*, increasing species complementarity among sites (Arita and Rodríguez, 2004, 2002; Balvanera et al., 2002; Sepkoski, 1988; Stendera and Johnson, 2005).

The reasoning presented above resembles the intermediate disturbance concept, where the highest species richness is found in sites with increased environmental disturbance (Connell, 1978; Fox, 1979; Huston, 1979). The difference between these two models lies in the temporal scale, once disturbance is defined as "a discrete event in time that changes the structure of an ecosystem, community, or population by changing resources, substrate availability, or physical environment". Adversity, in turn, would be "the high frequency of population reduction or conditions that result in a reduction in population growth rates" (Huston, 1979), and can be considered as an uninterrupted series of disturbances and can be regarded as an uninterrupted series of disturbances. Thus, the adversity variation spectrum would comprise the intensity of disturbance rather than its frequency, with a more coherent frequency leading to a reduced richness and abundance of communities in lotic environments (Death and Winterbourn, 1995).

Two problems are found in the intermediate disturbance model, where the first is the positive linear relationship between primary productivity and local diversity, and the second the fact that organism mobility reduces competitive exclusion, making it difficult to understand and test this hypothesis (Death, 2002). These very problems would also occur in the model proposed here, since it is an extrapolation of the intermediate disturbance hypothesis. Given the lack of information on the biology of benthic organisms, we decided to continue the thought process with these static processes, so that we could work only with integrity variation. However, understanding this interference can help to comprehend the observed pattern.

The difference between the areas of the estimated richness curves related to the relevance of *d* will then represent the change in species alternation, when related to a reduction in site integrity. Thus, the analytical difference is calculated by:

(2) $\frac{d_{\beta}}{d_{Ha}}=\sum o\left( s_{i} \right)-\sum o\left( s_{i} \right)\phi_{i}$ ,

where $\frac{d_{\beta}}{d_{Ha}}$ is the diversity turnover in respect to habitat integrity variation (*Ha*), the first summation is the potential richness and the second is the local richness because the *φ_i_* represent the niche overlap between the species. However, there is an analytical problem, once species richness is given by a Poisson distribution according to the anthropic impact on the stream. Negative values cannot be modeled by this distribution, so extreme fluctuations in the *a priori* distributions of potential and observed richness may show unlikely values. To solve this problem, we can reorganize (2):

(3) $\sum o\left( s_{i} \right)=\sum o\left( s_{i} \right)\phi_{i}+\frac{d_{\beta}}{d_{Ha}}$ .

The sum of potential species, named here local species pool (LSP), can be modeled using the complementary probability of non-occurrence of individuals of each species in the specific local (Σ1 – e ^– μ^). This artifact allows accounting all the species that occurred in the local with at least one individual. In other hand, the sum of realized occurred species, named her local species richness (LSR), can be modeled with a poison distribution of the number of species in the local. The difference of LSP and LSR, the diversity turnover in respect habitat integration variation, we can use a beta distribution to modelling the pattern observed. We choice a beta distribution, because this distribution has flexibility to fit a several distinct patterns. Now, we can adjust the equation (3) in the new analytical solution:

(4) $\sum1-e^{-\mu}=Poiss\left( S_{x} \right)+Beta\left( Ha,\alpha,\beta\right)$.

This equation measures the punctual richness turnover using the static habitat integrity value. Given that we are interested in the change of composition in an environmental gradient, *Ha* becomes a continuous parameter (between 0 and 1 following the HII value). These parameters of beta distribution change the local mean of individuals of each species, and we need modeling the parameter μ for all species related to environmental gradient. So, when we put the environmental gradient in the (4), the results is:

(5) $\sum1-e^{-Beta\left( x_{j},a_{i},b_{i} \right)}=Poiss\left( S_{x_{j}} \right)+Beta\left( x_{j},\alpha,\beta\right)$ ,

where *a*_i_ and *b*_i_ are the parameters of the beta distribution for genus *i*, given that each taxa responds differently to the environmental gradient, and *x*_j_ is the HII value in the stream *j*.

Computationally, the algorithm we used in R was:

Model{

for(j in 1:M){

a[j]~dgamma(0.001,0.001)

b[j]~dgamma(0.001,0.001)

for(i in 1:N){

mu[i,j]<-pow(x[i],a[j]-1)*pow(1-x[i],b[j]-1)*exp(loggam(a[j]+b[j])-

loggam(a[j])-loggam(b[j]))

p[i,j]<-1-exp(-mu[i,j])

muC[i,j]<-mu[i,j]+delta[i]

y[i,j]~dpois(muC[i,j])

pC[i,j]<-1-exp(-muC[i,j])}}

for(i in 1:N){

delta[i]<-pow(x[i],alfa-1)*pow(1-x[i],beta-

1)*exp(loggam(alfa+beta)-loggam(alfa)-loggam(beta))

somaC[i]<-sum(pC[i,])

soma[i]<-sum(p[i,])

riqueza[i]~dpois(soma[i])}

alfa~dgamma(0.001,0.001)

beta~dgamma(0.001,0.001)}

**Appendix B**

Model validation for Bayesian analysis

The results for *α* and *β* in the model, using gamma or uniform *priori* distribution in the models (Figure A.2). The bar represents the 95% credible interval.

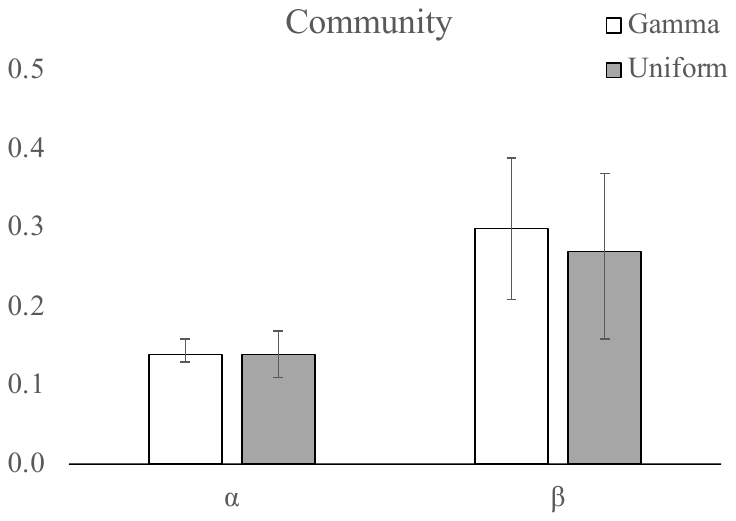

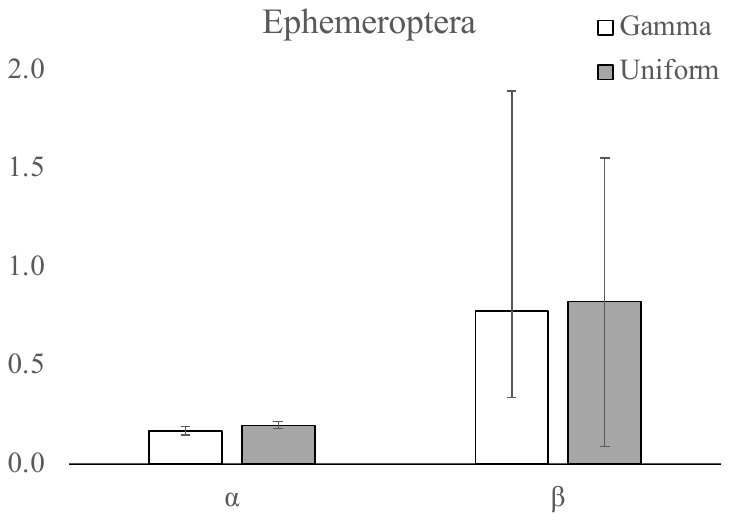

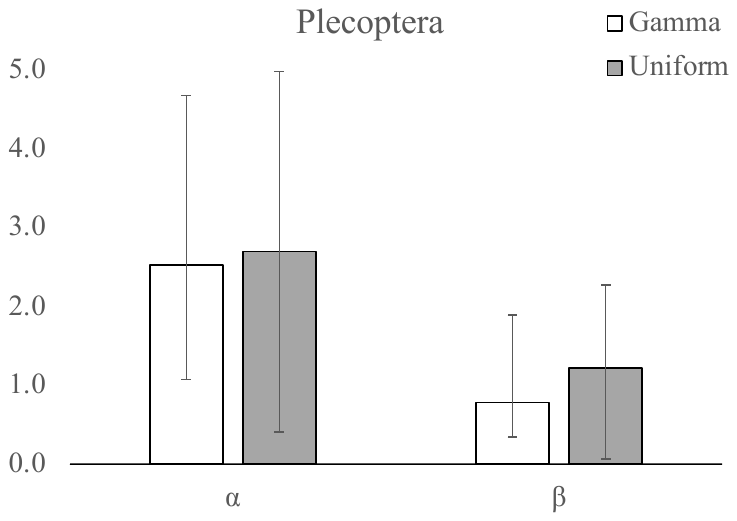

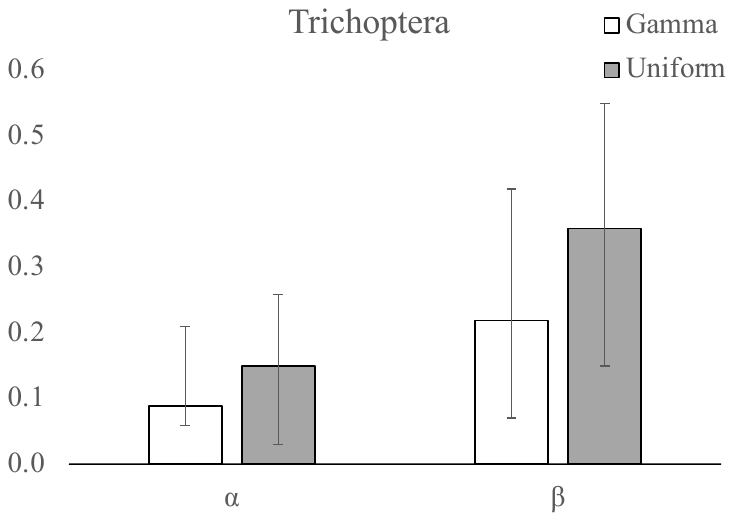

Figure A.2. The *posteriori* distribution for *α* and *β* with distinct *a priori* distribution for all community, Ephmeroptera, Plecoptera and Trichptera order.
